# Human.miRFFL.DB-A curated resource for human miRNA coregulatory networks and associated regulatory-circuits

**DOI:** 10.1101/2020.05.16.097865

**Authors:** Pankaj Khurana, Rajeev Varshney, R Sugadev, YK Sharma

**Affiliations:** Defence Institute of Physiology and Allied Sciences (DIPAS), Defence R&D Organization (DRDO), Timarpur, Delhi-110054

**Keywords:** miRNA:TF:TG coregulatory network, regulatory-circuits, miRNA-target interactions, Regulatory Networks

## Abstract

Gene regulation is viewed as a complex process where regulatory elements and their targets form highly complex network interactions thus affecting normal biological physiology and also disease-initiation and progress. Transcription factors (TF) and microRNA (miRNA) are fundamental transcriptional and post-transcriptional regulators of the gene expression controlling important biological processes. In recent years, many high throughput studies revealed that the complex regulatory interactions are mediated by the complex interplay between miRNA and TF regulating a Target Gene (TG) in conjunction. miRNAs and TFs are also known to regulate each other. This complex coregulatory mechanism may be represented in the form of miRNA:TF:TG coregulatory network. This network can be used to identify several small recurring subgraphs known as regulatory-circuits. One of these regulatory-circuits also called the Feed-Forward Loops (FFLs) is a three-node pattern which is composed of a miRNA and a TF, one of which regulates the other and both jointly regulate a TG. These regulatory-circuits have proven useful in elucidating the complex interplay of gene regulation during many physiological and pathological conditions.

Human.miRFFL.DB is a comprehensive integrated resource for human miRNA:TF:TG coregulatory directed networks and their associated regulatory-circuits. In-house scripts based on the graph theory principle have been used to identify both types of FFL motifs i.e. miRNA-FFL and TF-FFL. The database additionally provides an interactive visualization of the coregulatory networks and associated FFLs. Human.miRFFL.DB can be used as a comprehensive ready reference for human miRNA:TF:TG coregulatory networks and associated FFLs for decrypting complex cellular interactions of these regulatory biomolecules. Human.miRFFL.DB is available online at http://mirffldb.in/human/

## 1. Introduction

Gene expression regulation is a complex process involving various regulatory biomolecules across numerous levels [1]. Transcription factors (TFs) and microRNAs (miRNAs) are the two most common regulatory biomolecules that fine-tune gene expression by regulating at transcriptional level and post-transcriptional level respectively [2]. They are known to regulate gene expression independently; but recent increasing evidence show that miRNAs and TFs also work synergistically in the form of complex networks to regulate the gene expression, which further modulates cellular and molecular processes [3]. Their combinatorial role in disease initiation, processes, and recurrence has also been studied [4–6]. These complex regulatory interactions can be best studied and analysed using miRNA:TF:TG co-regulatory networks. These co-regulatory networks are responsible for the impressive degree of complexity in gene-regulation in higher eukaryots [7]. Studies on recurrent circuits, also known as network motifs, in gene co-regulatory network have significantly contributed to uncover this complexity, and in better understanding the regulatory roles of miRNAs and TFs [7]. Specifically, miRNAs and TFs have been shown to regulate common genes in these regulatory-circuits [8]. One of these regulatory-circuits called Feed Forward Loops (FFLs) are tripartite motifs in which miRNA regulates TF and/or regulated by it, and both together regulate the Target Gene (TG) in the network [9]. Based on type of interaction between miRNA and TF, the FFLs can be broadly classified into 2 types: TF-FFL and miRNA-FFL[10]. In TF-FFL, the TF regulates the miRNA and the TG while the miRNA represses the same common TG. In miRNA-FFL, the miRNAis the master regulator as it represses both the TF and TG while the TF also regulates the same TG (Figure 1). At the network level, both miRNA-FFLs and TF-FFLs are important genetic overrepresented motif patterns that occur more often than by chance in biological networks. Hence they serve as basic building blocks of a complex regulatory system [11, 12]. These FFLs are extensively studied to discover underlying genotypic and phenotypic relationship in complex disease conditions. e.g. TF: c-Myc; miR: miR-17-5p; target: E2F1 and c-Myc; miR: miR-20a; target: E2F1 were identified as module tightly regulating proliferative signal of proto-oncogene c- Myc and promoting cell cycle progression[13, 14]. C-Myc was termed as mater regulator of cellular proliferation in most of the human cancer types [15]. Hence c-Myc regulated FFL motifs in human cancer is extensively studied and a separate catalogue of C-myc regulating FFL motifs is consolidated [16]. Apart from the c-Myc other important FFLs regulating major molecular functions during cancer are TF: P53; miR: miR-34; target: MET, TF: E2F; miR: miR-106b/93/25; target: CDK, TF: AP-1; miR: miR-101; target: MMP9. These are known to regulate the extent of cancer invasion, anti-proliferative activity and cell migration respectively in different cancer studies [7]. A database of Cancer-Specific MicroRNA And Transcription Factor Co-Regulatory Networks (CMTCN) was recently developed to identify miRNA:TF:TG coregulatory network and associated FFLs in various human cancer types[17]. miRNA:TF:TG coregulatory networks and FFLs have also been found to be very useful in identifying mechanisms involved in other multifactorial disease for disease-initiation, -progression and -recurrence e.g. TF: MAX; miR: miR-320; target: BMP6 and TF: IRF1; miR: miR-103a-3p; target: ACVR2B was overrepresented in myocardial infarction recurrence condition and both targets BMP6, ACVR2B have been identified as biomarker for myocardial infarction recurrence[18]. Similarly TF: TP53; miR: mir-34b; target: CAV1, was demonstrated to be only dysregulated module in cardiac hypertrophy patients [19]. 9 FFLs, four miRNA-FFLs and five co-regulatory FFLs were found to be regulating cell-cycle regulation during a hypoxia stress[20].

**Figure 1:**
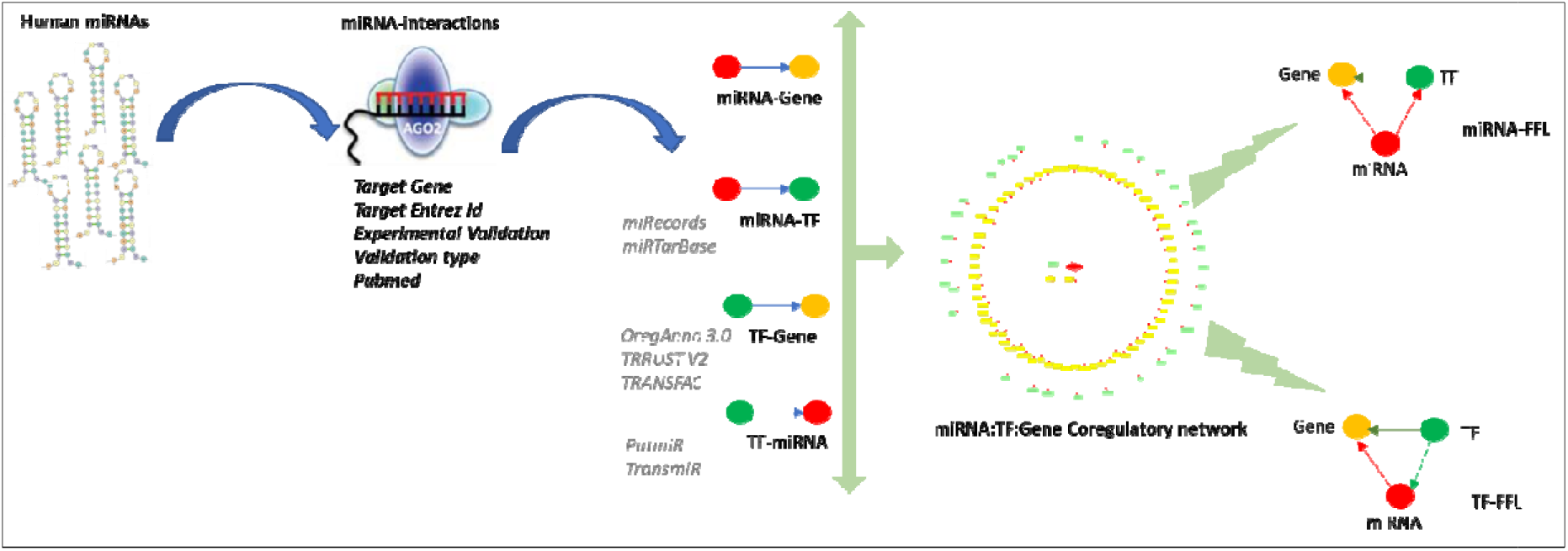
Overview of data collection and annotation in Human.miRFFL.DB database.

There are many resources that catalogue the miRNA:TF:TG coregulatory network and associated FFLs e.g. RegNetwork: regulatory network repository for gene regulatory networks (GRNs) provides different types of regulatory interactions i.e‘TF:TG’,‘TF:miRNA’, miRNA:TG’ ‘miRNA:TF’[21]. CircuitsDB: is a database that provides mixed miRNA/ TF-FFL circuits of 193 mature miRNAs and 130 pre-miRNAs of human and mouse[22]. TFmiR and MAGIA2 are webservers that uses miRNAs/mRNAs expression profile to identify significantly enriched TF and miRNAs interactions for construction of miRNA:TF:TG coregulatory network[23, 24]. Though there are several databases and tools available for FFL identification but these resources have certain limitations. Some database are not comprehensive, whereas others require an expression profile of miRNAs or TFs for identification of FFL. Hence, some FFLs which are biologically relevant may be lost during enrichment processes. Hence there is a vital need for a comprehensive global resource of human miRNA:TF:TG coregulatory networks and associated FFLs.

Here, we present Human.miRFFL.DB, a comprehensive user-friendly database of human miRNA: TF: TG coregulatory directed networks and their associated regulatory-circuits. The current version of Human.miRFFL.DB contains miRNA:TF:TG coregulatory networks of ~2600 human miRNAs. To construct miRNA:TF:TG coregulatory networks, Human.miRFFL.DB has an exhaustive list of 5,02,652 validated miRNA- Target Interactions (MTIs). The database stores experimentally validated MTIs with associated details like miRNA-target name, target Entrez ID, experimental validation type, and support experimental validation (Functional/Functionally Weak) and link to corresponding publication that are displayed in tabular format. Human.miRFFL.DB characterizes these MTIs as miRNA:TG or miRNA:TF interactions. TF:TG and TF:miRNA interactions are also added to construct miRNA:TF:TG directed coregulatory network. Human.miRFFL.DB offers interactive visualization of this network in which the nodes are color-coded for their easy characterization (TG as yellow, TF as green and miRNA as red). Human.miRFFL.DB uses in-house scripts based on graph theory principle to identify all tripartite miRNA-FFL and TF-FFL motifs from the network. It also provides an interactive visualization of each tripartite graph. The MTIs, miRNA-FFLs and TF-FFLs, can be downloaded in excel and PDF format. The network file can also be downloaded in .SIF format, which can be used with popular visualization software like Cytoscape, Gephi, BINA. Hence Human.miRFFL.DB can be considered as a comprehensive non-redundant catalogue of human miRNA:TF:TG coregulatory networks and its associated FFLs. We hope that Human.miRFFL.DB will catalyze research in understanding the complex crosstalk between miRNA, TF and TG. These motifs may offer mechanistic insights into the complex regulatory mechanisms. Combining it with experimental validation, these FFLs can identify novel players that can be used as diagnostic or prognostic markers and therapeutic targets for multifactorial complex disorders and pathophysiological conditions.

## 2. Material and Methods

### 2.1 Data Collection and Processing

To construct Human.miRFFL.DB database, a non-redundant list of human miRNA was retrieved from miRBase (version 22). To ensure uniformity in the nomenclature, the precursor miRNAs were mapped to their mature form using miRBase[25] and miRDB[26] repository. Dead/obsolete entries were removed. Experimentally validated target interactions of each human miRNA was fetched from public repositories i.e. mirTarBase[27] and miRecords[28] using multiMiR[29] R package. For each interaction, additional information like target name, target Entrez ID, support experiment details, support experimental validation (functional strong/functionally Weak) and the reference PMID was also fetched. Figure 1 schematically illustrates the general workflow for the collection of resource data for Human.miRFFL.DB.

### 2.2 Data Enrichment

The data was further enriched by labelling miRNA Target Intercations (MTIs) as miRNA: TG and miRNA:TF interaction by using the universal dataset of all human 3738 TFs and co-transcription factors (coTFs). A list of 3462, 3240, 1769, 1758, 3360, 1538 TFs were fetched from TFcheckpoint[30], DBD[31], ORFeome[32], TcoF-DB V2[33], TFCat[34], TFClass[35], TRANSFAC[36] databases respectively. A non-redundant list of 3292 unique TFs were obtained after removing duplicate entries. Further 958 coTFs were also identified from TcoF-DB V2 database [33]. Thus a combined list of 3798 unique TFs and coTFs was obtained. The TF: TG interactions were fetched from TRANSFAC, OregAnno 3.0 [37] and TRRUST V2[38] databases. TF:miRNA interactions were fetched from TransmiR[39] and PuTmiR[40] databases. Finally, all the dataset files were loaded as JavaScript Object Notation (JSON) files format and stored in the MongoDB database. Vis.js specifically was used to display miRNA: TF:TG networks.

### 2.3 FFL Motif Identification

Each vertex in the network is labelled as V_m_, V_TF_ or V_TG_. Where, V_m_ refers to a miRNA, V_TF_ refers to a TF and V_TG_ refers to a TG. The edges are annotated as E_mt,_ E_mg,_ E_tg,_ E_tm_ where E_mt_ refers to edge from V_m_ to V_TF_, E_mg_ refers to edge from V_m_ to V_TG_, E_tg_ refers to edge from V_TF_ to V_gene_ and E_tm_ refers to edge from V_TF_ to V_m._

For each Vm, its edges E_mg_ from the miRNA:TG interactions is searched in miRNA:TF interaction to identify a corresponding Emt with a common vertex Vm. Thereafter, an analogous Etg is searched in the TF:TG interactions. If found, the complete graph containing three vertices (Vm, V_TF_, V_TG_) and edges(E_mt,_ E_mg,_ E_tg_) were labelled as miRNA-FFL motif graph.

Similarly in order to find TF-FFL motif, edge E _*tm*_ and *E_tg_* were identified from TF-miRNA and miRNA:TG interactions dataset respectively. Further the program scans for an E_mg_ associated with V_m_ and V_TG_ in miRNA:TG interaction dataset. If algorithm finds E_mg_, the complete graph containing three vertices (Vm, V_TF_, V_TG_) and edges(E_tm,_ E_mg,_ E_tg_) were labelled as TF-FFL motif graph. This methodology was used to identify miRNA-FFLs and TF-FFLs for each human miRNA.

### 2.4 Database development

All database files were collected, processed and stored as JavaScript Object Notation (JSON) files in MongoDB database[41]. MongoDB is an open-source document-oriented (NoSQL) database [41, 42]. The variables and query functions of the application program interface (API) in the Human.miRFFL.DB web application are defined in the python language. The database uses Asynchronous JavaScript and XML (AJAX) technique for API calls[43]. AJAX is a web development technique that is used for creating interactive web applications. It utilizes XHTML for content along with the document object model and JavaScript for dynamic content display. Vis.js library functions are used for interactive visualization of network graphs on front-end.

## 3. RESULTS

Human.miRFFL.DB is a comprehensive non-redundant resource for human miRNA:TF:TG coregulatory directed networks and their associated regulatory-circuits. It contains a non-redundant list of 2596 human miRNAs, 11,864 experimentally validated TGs and 3,798 TFs including co-Transcription cofactors (coTFs) (Figure 2a). The database contains ~5,00,000 experimentally validated MTIs mined from mirTarBase[27], miRecords[28] and ~7200 corresponding reference PMIDs. The ~3,80,000 unique miRNA-target interactions are categorized as ~1, 14,000 miRNA: TF interactions and ~2, 66,000 miRNA: TG interactions.

**Figure 2.**
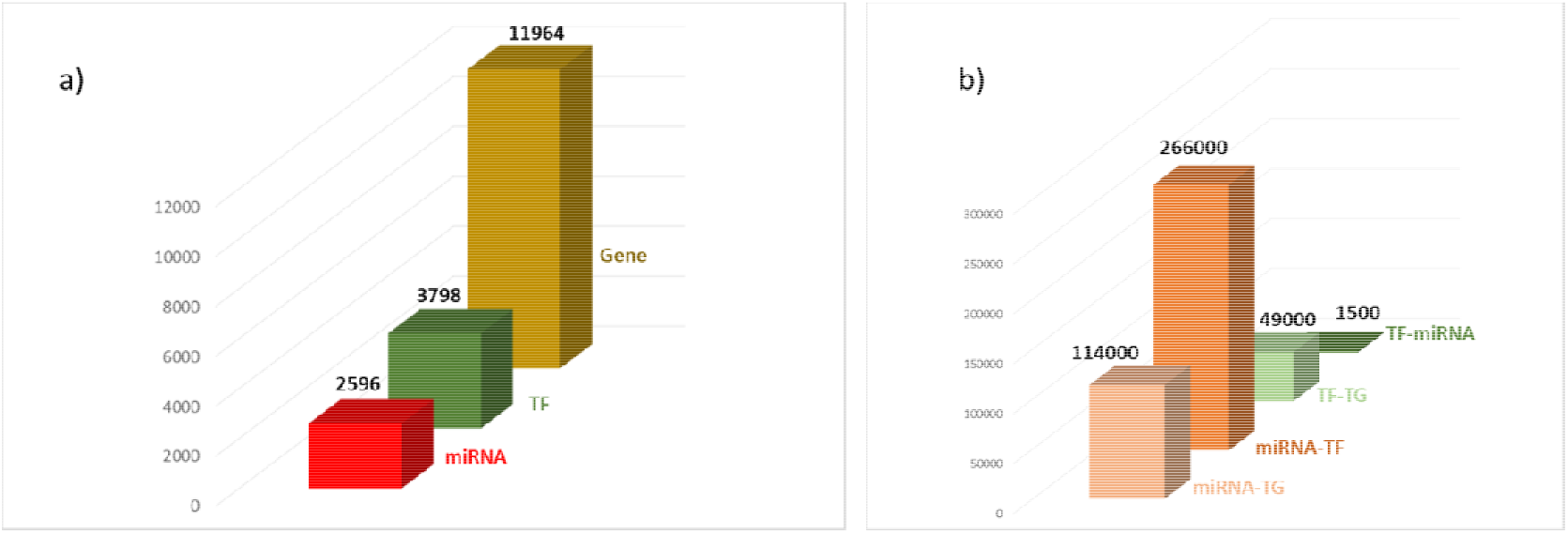
a) Number of miRNAs, TFs and TGs in Human.miRFFL.DB. b) Number of miRNA:TG, miRNA:TF, TF:TG, TF:miRNA interactions in the database.

**Figure 3:**
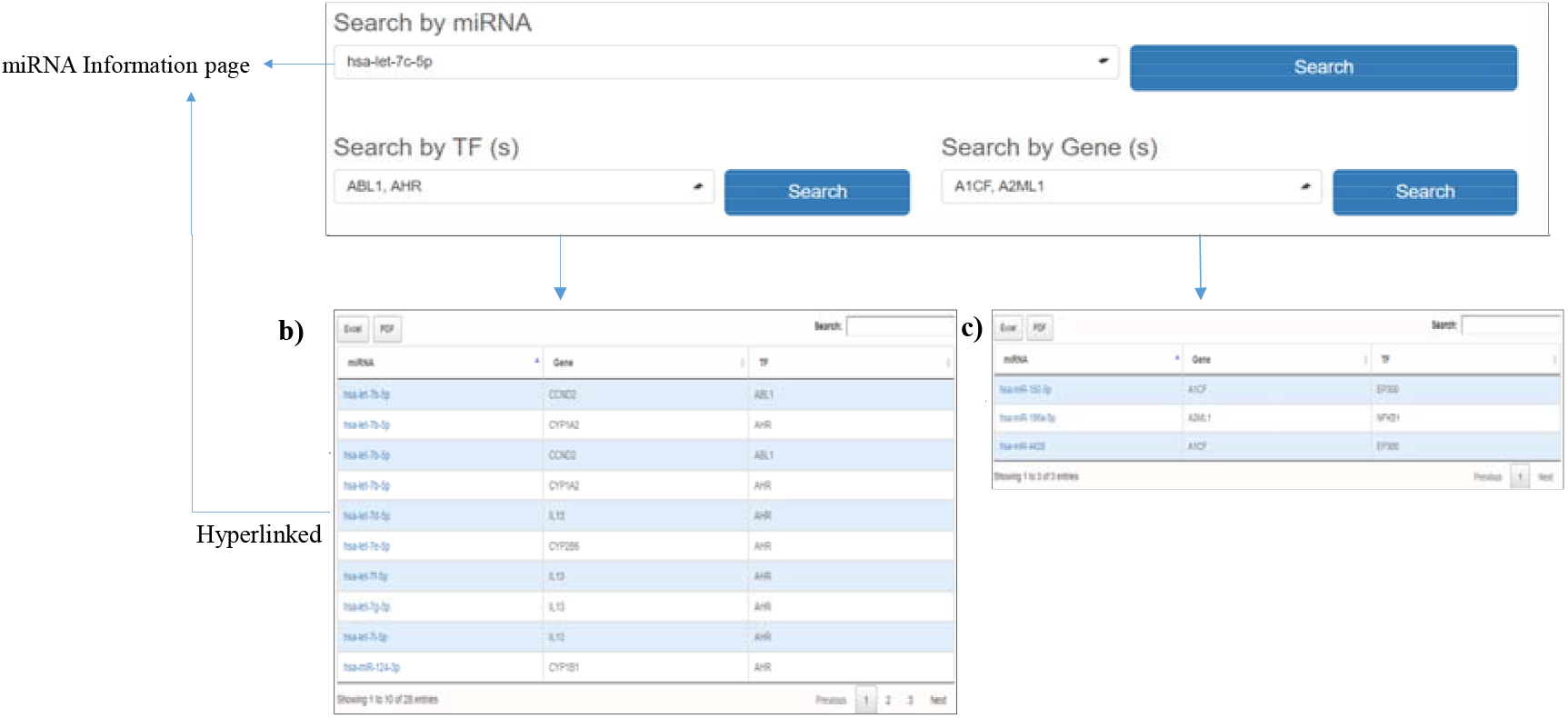
The figure shows the screenshot of the ‘Browse’ option. This option allow the users to browse miRNAs using three different options: ‘Search by miRNA’, ‘Search by TF(s)’ and ‘Search by Gene(s)’.a) On searching with miRNA, the page connects to the detailed information page of the respective human miRNA. b) Table representing the FFLs containing the queried TF(s) c) Table representing the FFLs containing the queried gene(s).

Additionally, ~49,000 TF:TG interactions were fetched from OregAnno[37] and TRRUST V2[38] databases and 1500 TF:miRNA interactions were fetched from TransmiR[39] and PuTmiR[40] (Figure 2b). These interactions were used to construct miRNA:TF:TG coregulatory directed network for each human miRNA.

### 3.1 Web Interface

Human.miRFFL.DB offers browsing by three routes i.e. ‘search by miRNA’, ‘search by TF(s)’ or ‘search by gene(s)’ (Figure 2). ‘Search by miRNA’ allows the user to select miRNA from a pull down menu. On clicking the search button, the page connects to the detailed information page of the respective miRNA, which is discussed in detail in next section 2.2. The database also facilitates the user to search the database by selecting TF(s) and TG(s) from a pull down menu. On searching the user can view the FFLs containing the queried TF(s) or gene(s) in a tabular format (Figure 2b, 2c). The table can be downloaded in Excel and PDF format. The miRNAs of the FFLs are further hyperlinked to the corresponding information page.

### 3.2 miRNA Information Page

The miRNA information page for each human miRNA can be divided into two sections. The first section displays interactive visualization of miRNA:TF:TG coregulatory network and a tabular representation of miRNA-FFLs and TF-FFLs that are identified from the network. The second section enlists the experimentally-validated MTIs and its associated information in a tabular format.

#### 3.2.1 miRNA:TF:TG co-regulatory network and FFLs

The top of the miRNA information page highlights the name of the mature miRNA which is hyperlinked to miRBase database[25]. miRBase database is a comprehensive resource of miRNA sequence data, annotation and predicted gene targets. Hence hyperlinking each human miRNA in the database serve as a ready reference to get additional details about these miRNAs that includes mature and precursor miRNA sequence, miRNA stem loop structure, chromosome location, source organism etc (Figure 4).

**Figure 4:**
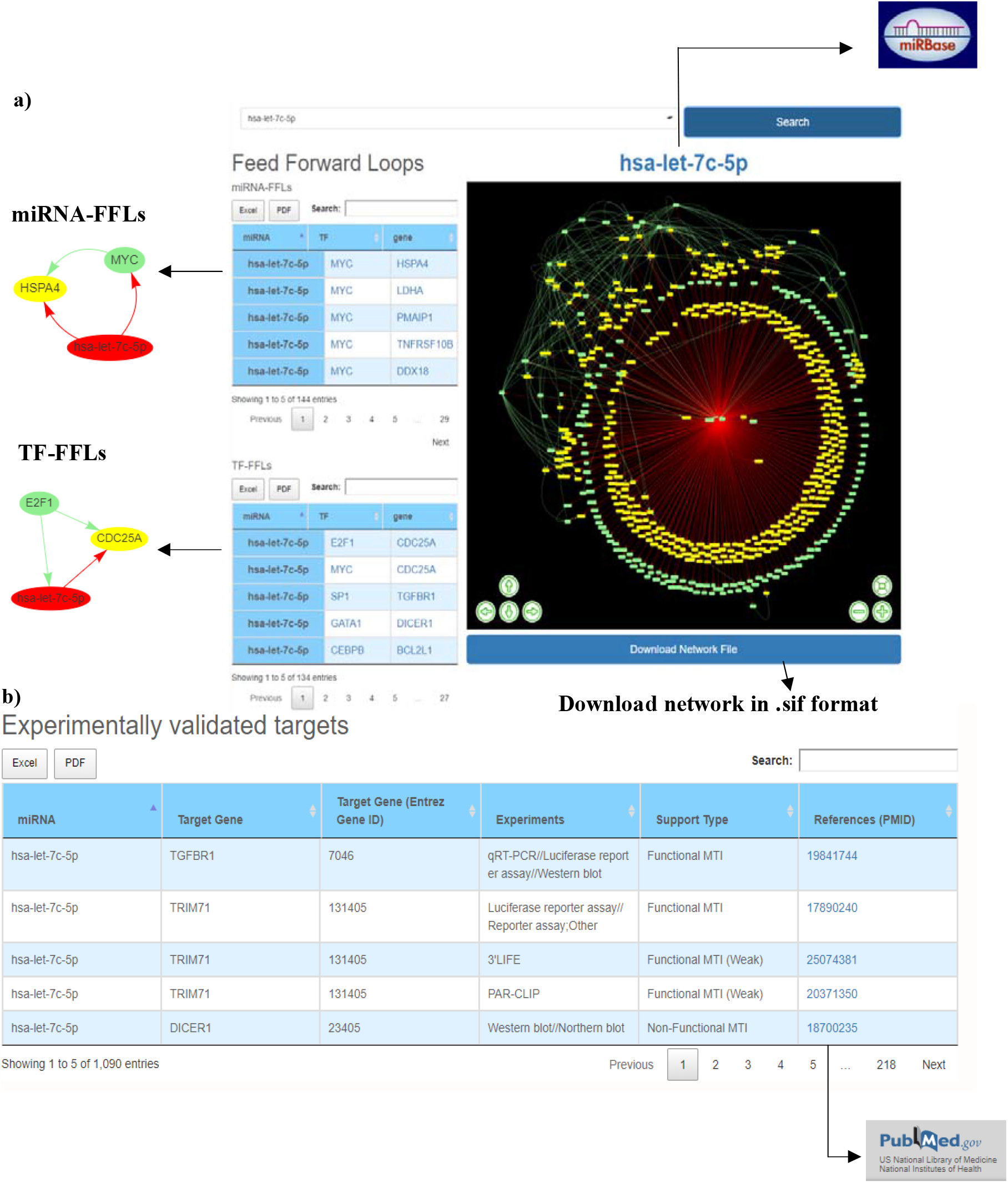
The web image of miRNA information page of hsa-let-7c-5p. a) The first section shows the coregulatory network and the associated miRNA-FFLs and TF-FFLs. Clicking on the miRNA, a pop-up window displays the corresponding FFL. The gene names are hyperlinked to GeneCards database, which provides additional details about the gene. b) The experimentally validated targets of the miRNA are listed along with the details like Target gene entrez ID, experiment, support type and the respective reference paper. The PMID is hyperlinked to the PubMed database for ready access to the publication.

The miRNA:TF:TG network is displayed as an interactive network in which the nodes are color coded for easy characterization (miRNA in red and its interacting TFs and TGs in green and yellow respectively) (Figure 4a). The network is also equipped with zoom-in, zoom-out, and node-translation options. The network file can be downloaded in (.SIF) format and can be readily used by many open-source software’s such as Cytoscape[44], Gephi[45], BINA[46], etc for further analysis.

The miRNA-FFL and TF-FFL are identified from the coregulatory network using in-house scripts and presented as separate tables. Each row of the table represents a FFL with a miRNA, TF and TG. User can also visualize tripartite graph by clicking on the miRNA, which will appear in a pop-up window (Figure 4a). Both the miRNA-FFL and TF- FFL tables are provided with an integrated search option. This allows the user to search the lengthy tables by TF/gene symbol. Each TF and TG symbol in the FFL table are further hyperlinked to the GeneCards[47] database. This would serve as a ready reference for the user to get additional details about the gene that includes aliases of genes, promoter and enhancer location of gene, protein coded by gene, functional characterization, cellular localization, pathway enrichment, gene-gene interaction network, drug-gene relationship, tissue specific gene expression profile, orthologs, paralogs, transcript variant etc.

#### 3.2.2. Experimentally validated MTIs

Human.miRFFL.DB also provide experimentally validated MTIs along with its associated information that includes miRNA-target symbol, miRNA-target Entrez ID, experimental validation type, and support experimental validation (Non-Functional/Functional/Functionally Weak) and the reference paper. The information is provided in a tabular format that can be downloaded in the form of excel/PDF file. The table is also provided with an integrated search option, which allows to search the lengthy tables. The user may search with target symbol, entrez ID, experiment type, support type or PMID (Figure 4b). The PMID is hyperlinked to the PubMed database for ready access to the publication.

## 4. Discussion

A decade earlier, Yitzhak Pilpel et al studied the global miRNA:TF:TG mammalian co-regulatory network and uncovered two network architecture of regulatory network [48]. The network analysis of this global miRNA:TF mammalian co-regulatory network revealed several recurring motifs of miRNA and TF regulating a hub gene. These motifs or regulatory-circuits help in fine-tuning of many complex molecular and cellular processes. Since then, studies on recurrent circuits in regulatory networks, also known as network motifs, have substantially contributed to addressing this complexity and in particular to better understand miRNAs and TFs exert their regulatory roles[5, 7, 9]. The most studied recurring motifs in miRNA:TF:TG coregulatory networks are FFLs. FFLs have been proven to be the best network analysis tool to study the combinatorial TG regulation by miRNA and TF in many complex pathologies and are crucial in providing new insights into the logic and evolution of a new regulatory layer of the complex eukaryotic genome. Human.miRFFL.DB offers a comprehensive and interactive collection of human miRNA:TF:TG coregulatory networks and associated regulatory-circuits. Human.miRFFL.DB pools miRNA:TG, miRNA:TF, TF:TG and TF:miRNA interactions for constructing complex miRNA:TF:TG coregulatory networks of human miRNAs. These networks were used to identify the miRNA-FFL and TF-FFL motifs. Additionally, Human.miRFFL.DB also provides experimentally validated miRNA-Target Interactions (MTIs) resources for these human miRNAs.

A thorough literature survey suggested that though there are several server platforms, databases, and tools available for identification of gene regulatory networks (GRNs) and associated FFLs; all these research tools lack in addressing the core of the problem according to the current scenario e.g. CircuitsDB: a database for mixed miRNA/ TF FFL circuits in human and mouse contains information of only 193 mature miRNAs and 130 pre-miRNAs FFLs. Whereas, currently more than ~2600 human mature miRNAs (miRBase release 22) and ~ 5,00,000 experimentally validated MTIs have already been identified[22, 27, 49]. RegNetwork: an integrated resource of transcriptional and post-transcriptional regulatory networks provides a collection of regulatory interactions among TFs, miRNAs and target genes. [21]. The database, however, does not provide FFLs. Also in the thorough examination of the database, many interactions were found missing. A random example of hsa-miR-141-3p has been used to compare the regulatory interactions present in our database (Table S1). Server tools such as TFmiR, MAGIA2 require miRNAs/mRNAs expression profile as input to identify regulatory interactions based on the input deregulated genes and deregulated miRNAs [23, 24]. For identification of regulatory interactions, they first identify miRNAs, whose target genes, as well as TFs, are significantly enriched within the input deregulated genes by using the hypergeometric distribution function. Hence only interactions among these miRNAs and significantly enriched TF, TGs are used to construct miRNA:TF:TG coregulatory network [23, 24]. This methodology reduces a significant amount of interactions from the network. Subsequently, it would also lead to the loss of important FFLs identified from the network. Databases like ‘CMTCN’ [17] and ‘curated database of miRNA mediated feed-forward loops including MYC as master regulator’[16] provide only the cancer-specific miRNA:TF:TG coregulatory networks and associated FFLs. Though these databases are recently published their global coverage is limited to only cancer. FFLs have also been identified in many other multifactorial diseases and pathophysiological conditions like myocardial infarct, schizophrenia, and hypoxia respectively. So these databases, webservers, and tools cannot be considered universal repository for miRNA:TF:TG coregulatory networks and FFLs. Herein, we developed Human.miRFFL.DB which is a comprehensive coregulatory network resource that integrates miRNA:TG interactions, miRNA:TF interactions, TF:TG interactions and TF: miRNA interactions to build co-regulatory networks. These complex networks provide a global representation of complex regulatory and target interactions of human miRNAs and TFs. It can advance our understanding of complex molecular mechanisms that are controlled by these regulatory molecules.

## 5. Conclusion

Human.miRFFL.DB identifies FFLs that will help to decode the interplay between miRNAs, TFs and TGs to get new mechanistic-insights into specific molecular and cellular processes. Further analysis of these FFL motifs using network biology techniques could also help us in identifying potential disease-markers and therapeutic targets.The resource can also be used for other network-based computational and integrated analysis. This would allow researchers to be more focused on analysis rather than going for manual curation of the interactions from different databases to construct these co-regulatory networks. Also it is ready reference of enriched FFLs for human miRNA pool. We believe that Human.miRFFL.DB will help researchers to design and also analyze miRNA based experiments.

## Supplementary Materials

Table S1: Tabular representation of comparison in regulatory interactions of the hsa-miR-141-5p miRNA present in RegNetwork database with Human.miRFFL.DB.

## Abbreviations

TF: Transcription factor
TG: Target Gene
FFL: Feed Forward Loop
MTI: miRNA-Target Interaction
coTF: Co Transcription Factor

