## Supplementary material for "Human.miRFFL.DB-A curated resource for human miRNA coregulatory networks and associated regulatory-circuits": Table S1

Table S1: Shows comparison of regulatory interaction data of miRFFLDB database and RegNetwork database using example search of hsa-miR-141-3p.

| **miRFFLDB** | | | **RegNetwork** | | |
| --- | --- | --- | --- | --- | --- |
| **Regulator symbol** | **Target symbol** | **Experimental validation** | **Regulator symbol** | **Target symbol** | **Evidence** |
| hsa-miR-141-3p | QKI | validated | E2F1 | hsa-miR-141-3p | Predicted |
| hsa-miR-141-3p | CHML | validated | EGR1 | hsa-miR-141-3p | Predicted |
| hsa-miR-141-3p | PTPRD | validated | FOS | hsa-miR-141-3p | Predicted |
| hsa-miR-141-3p | GNA13 | validated | GABPA | hsa-miR-141-3p | Predicted |
| hsa-miR-141-3p | STXBP2 | validated | JUN | hsa-miR-141-3p | Predicted |
| hsa-miR-141-3p | CELF1 | validated | MAX | hsa-miR-141-3p | Predicted |
| hsa-miR-141-3p | CDV3 | validated | MXI1::CLEC5A | hsa-miR-141-3p | Predicted |
| hsa-miR-141-3p | CDC25C | validated | MYC | hsa-miR-141-3p | Predicted |
| hsa-miR-141-3p | CDC25A | validated | NFYA | hsa-miR-141-3p | Predicted |
| hsa-miR-141-3p | PRELID2 | validated | PAX5 | hsa-miR-141-3p | Predicted |
| hsa-miR-141-3p | MCL1 | validated | PPARG | hsa-miR-141-3p | Predicted |
| hsa-miR-141-3p | PPP1R15B | validated | SP1 | hsa-miR-141-3p | Predicted |
| hsa-miR-141-3p | SPRYD4 | validated | SPI1 | hsa-miR-141-3p | Predicted |
| hsa-miR-141-3p | FUT11 | validated | TAL1 | hsa-miR-141-3p | Predicted |
| hsa-miR-141-3p | ZMPSTE24 | validated | TCF3 | hsa-miR-141-3p | Predicted |
| hsa-miR-141-3p | MAP4K4 | validated | TFAP2A | hsa-miR-141-3p | Predicted |
| hsa-miR-141-3p | CCDC71L | validated | TFAP2C | hsa-miR-141-3p | Predicted |
| hsa-miR-141-3p | CCDC18 | validated | USF1 | hsa-miR-141-3p | Predicted |
| hsa-miR-141-3p | MALAT1 | validated |  |  |  |
| hsa-miR-141-3p | YWHAG | validated |  |  |  |
| hsa-miR-141-3p | YRDC | validated |  |  |  |
| hsa-miR-141-3p | YME1L1 | validated |  |  |  |
| hsa-miR-141-3p | XPOT | validated |  |  |  |
| hsa-miR-141-3p | PIGW | validated |  |  |  |
| hsa-miR-141-3p | PHLPP2 | validated |  |  |  |
| hsa-miR-141-3p | PHLPP1 | validated |  |  |  |
| hsa-miR-141-3p | SLC35D1 | validated |  |  |  |
| hsa-miR-141-3p | WDR37 | validated |  |  |  |
| hsa-miR-141-3p | C18orf25 | validated |  |  |  |
| hsa-miR-141-3p | C16orf58 | validated |  |  |  |
| hsa-miR-141-3p | PGK1 | validated |  |  |  |
| hsa-miR-141-3p | PGAM4 | validated |  |  |  |
| hsa-miR-141-3p | PEX11A | validated |  |  |  |
| hsa-miR-141-3p | VAC14 | validated |  |  |  |
| hsa-miR-141-3p | ERO1A | validated |  |  |  |
| hsa-miR-141-3p | KLHL20 | validated |  |  |  |
| hsa-miR-141-3p | USP53 | validated |  |  |  |
| hsa-miR-141-3p | EPHA7 | validated |  |  |  |
| hsa-miR-141-3p | EPHA2 | validated |  |  |  |
| hsa-miR-141-3p | BICRAL | validated |  |  |  |
| hsa-miR-141-3p | UQCRFS1 | validated |  |  |  |
| hsa-miR-141-3p | SHC1 | validated |  |  |  |
| hsa-miR-141-3p | KIAA1549L | validated |  |  |  |
| hsa-miR-141-3p | KIAA1147 | validated |  |  |  |
| hsa-miR-141-3p | PAPPA | validated |  |  |  |
| hsa-miR-141-3p | ELMO2 | validated |  |  |  |
| hsa-miR-141-3p | BAP1 | validated |  |  |  |
| hsa-miR-141-3p | EIF4E | validated |  |  |  |
| hsa-miR-141-3p | UBAP1 | validated |  |  |  |
| hsa-miR-141-3p | ATXN7L1 | validated |  |  |  |
| hsa-miR-141-3p | TROVE2 | validated |  |  |  |
| hsa-miR-141-3p | IPO5 | validated |  |  |  |
| hsa-miR-141-3p | TRMT112 | validated |  |  |  |
| hsa-miR-141-3p | DPY19L1 | validated |  |  |  |
| hsa-miR-141-3p | ARL5B | validated |  |  |  |
| hsa-miR-141-3p | DNAJC28 | validated |  |  |  |
| hsa-miR-141-3p | S100A6 | validated |  |  |  |
| hsa-miR-141-3p | TRAPPC2B | validated |  |  |  |
| hsa-miR-141-3p | TRAM1 | validated |  |  |  |
| hsa-miR-141-3p | IGF1R | validated |  |  |  |
| hsa-miR-141-3p | APOOL | validated |  |  |  |
| hsa-miR-141-3p | HSPA4 | validated |  |  |  |
| hsa-miR-141-3p | TNKS2 | validated |  |  |  |
| hsa-miR-141-3p | RPL12 | validated |  |  |  |
| hsa-miR-141-3p | TNFRSF10B | validated |  |  |  |
| hsa-miR-141-3p | ANGPTL7 | validated |  |  |  |
| hsa-miR-141-3p | TMOD3 | validated |  |  |  |
| hsa-miR-141-3p | HNRNPF | validated |  |  |  |
| hsa-miR-141-3p | NDST4 | validated |  |  |  |
| hsa-miR-141-3p | CYP1B1 | validated |  |  |  |
| hsa-miR-141-3p | RIN2 | validated |  |  |  |
| hsa-miR-141-3p | TM4SF1 | validated |  |  |  |
| hsa-miR-141-3p | ACVR2B | validated |  |  |  |
| hsa-miR-141-3p | TIAM1 | validated |  |  |  |
| hsa-miR-141-3p | COX6B1 | validated |  |  |  |
| hsa-miR-141-3p | RASSF2 | validated |  |  |  |
| hsa-miR-141-3p | RAP2C | validated |  |  |  |
| hsa-miR-141-3p | TGFB2 | validated |  |  |  |
| hsa-miR-141-3p | RAB8B | validated |  |  |  |
| hsa-miR-141-3p | CLDND1 | validated |  |  |  |
| hsa-miR-141-3p | A1BG | validated |  |  |  |
| hsa-miR-141-3p | TAZ | validated |  |  |  |
| hsa-miR-141-3p | KIAA1549 | validated |  |  |  |
| hsa-miR-141-3p | ELAVL4 | validated |  |  |  |
| hsa-miR-141-3p | KEAP1 | validated |  |  |  |
| hsa-miR-141-3p | ZNF621 | validated |  |  |  |
| hsa-miR-141-3p | ELAVL2 | validated |  |  |  |
| hsa-miR-141-3p | SFPQ | validated |  |  |  |
| hsa-miR-141-3p | PTEN | validated |  |  |  |
| hsa-miR-141-3p | PSMD11 | validated |  |  |  |
| hsa-miR-141-3p | CDYL | validated |  |  |  |
| hsa-miR-141-3p | E2F3 | validated |  |  |  |
| hsa-miR-141-3p | OGT | validated |  |  |  |
| hsa-miR-141-3p | STK3 | validated |  |  |  |
| hsa-miR-141-3p | STAT5A | validated |  |  |  |
| hsa-miR-141-3p | STAT4 | validated |  |  |  |
| hsa-miR-141-3p | IRF2BPL | validated |  |  |  |
| hsa-miR-141-3p | SCD5 | validated |  |  |  |
| hsa-miR-141-3p | MED13 | validated |  |  |  |
| hsa-miR-141-3p | MDM4 | validated |  |  |  |
| hsa-miR-141-3p | ZNF292 | validated |  |  |  |
| hsa-miR-141-3p | PRKAA2 | validated |  |  |  |
| hsa-miR-141-3p | GATA6 | validated |  |  |  |
| hsa-miR-141-3p | NR0B2 | validated |  |  |  |
| hsa-miR-141-3p | DLX5 | validated |  |  |  |
| hsa-miR-141-3p | MAPK9 | validated |  |  |  |
| hsa-miR-141-3p | ZMAT3 | validated |  |  |  |
| hsa-miR-141-3p | MAPK14 | validated |  |  |  |
| hsa-miR-141-3p | ZFPM2 | validated |  |  |  |
| hsa-miR-141-3p | PPARA | validated |  |  |  |
| hsa-miR-141-3p | ZEB2 | validated |  |  |  |
| hsa-miR-141-3p | ZEB1 | validated |  |  |  |
| hsa-miR-141-3p | POLR3F | validated |  |  |  |
| hsa-miR-141-3p | MALT1 | validated |  |  |  |
| hsa-miR-141-3p | MACC1 | validated |  |  |  |
| hsa-miR-141-3p | ZBTB34 | validated |  |  |  |
| hsa-miR-141-3p | HOXB5 | validated |  |  |  |
| hsa-miR-141-3p | HNRNPD | validated |  |  |  |
| hsa-miR-141-3p | HNRNPAB | validated |  |  |  |
| hsa-miR-141-3p | LPP | validated |  |  |  |
| hsa-miR-141-3p | YAP1 | validated |  |  |  |
| hsa-miR-141-3p | CTBP2 | validated |  |  |  |
| hsa-miR-141-3p | LHX1 | validated |  |  |  |
| hsa-miR-141-3p | HIPK2 | validated |  |  |  |
| hsa-miR-141-3p | ADNP2 | validated |  |  |  |
| hsa-miR-141-3p | PHB2 | validated |  |  |  |
| hsa-miR-141-3p | RBM28 | validated |  |  |  |
| hsa-miR-141-3p | HDGF | validated |  |  |  |
| hsa-miR-141-3p | RB1 | validated |  |  |  |
| hsa-miR-141-3p | H2AFZ | validated |  |  |  |
| hsa-miR-141-3p | TFDP2 | validated |  |  |  |
| hsa-miR-141-3p | BRD3 | validated |  |  |  |
| hsa-miR-141-3p | TET3 | validated |  |  |  |
| hsa-miR-141-3p | TET1 | validated |  |  |  |
| hsa-miR-141-3p | ZNF805 | validated |  |  |  |
| hsa-miR-141-3p | ERBIN | validated |  |  |  |
| hsa-miR-141-3p | KLF5 | validated |  |  |  |
| hsa-miR-141-3p | TCF7L1 | validated |  |  |  |
| hsa-miR-141-3p | KLF12 | validated |  |  |  |
| hsa-miR-141-3p | KLF11 | validated |  |  |  |
| hsa-miR-141-3p | BICD2 | validated |  |  |  |
| hsa-miR-141-3p | CLOCK | validated |  |  |  |
| ZEB1 | CYP1B1 | validated |  |  |  |
| MYC | EIF4E | validated |  |  |  |
| MYC | hsa-miR-141-3p | validated |  |  |  |
| EP300 | RASSF2 | validated |  |  |  |
| EP300 | hsa-miR-141-3p | validated |  |  |  |
| MYC | CDC25C | validated |  |  |  |
| EP300 | TM4SF1 | validated |  |  |  |
| MYC | CDC25A | validated |  |  |  |
| EP300 | PPP1R15B | validated |  |  |  |
| EP300 | RAB8B | validated |  |  |  |
| EP300 | QKI | validated |  |  |  |
| MYC | QKI | validated |  |  |  |
| MYC | SPRYD4 | validated |  |  |  |
| EP300 | PGK1 | validated |  |  |  |
| MYC | PGK1 | validated |  |  |  |
| EP300 | IPO5 | validated |  |  |  |
| EP300 | CYP1B1 | validated |  |  |  |
| KAT2B | CYP1B1 | validated |  |  |  |
| KAT2B | hsa-miR-141-3p | validated |  |  |  |
| ZEB1 | hsa-miR-141-3p | validated |  |  |  |
| EP300 | USP53 | validated |  |  |  |
| EP300 | HNRNPF | validated |  |  |  |
| MYC | HNRNPF | validated |  |  |  |
| MYC | PIGW | validated |  |  |  |
| MYC | XPOT | validated |  |  |  |
| EP300 | RPL12 | validated |  |  |  |
| MYC | RPL12 | validated |  |  |  |
| EP300 | TROVE2 | validated |  |  |  |
| MYC | HSPA4 | validated |  |  |  |
| EP300 | MALAT1 | validated |  |  |  |
| MYC | MALAT1 | validated |  |  |  |
| MYC | TNFRSF10B | validated |  |  |  |
